# A population code for idiothetic representations in the hippocampal-septal circuit

**DOI:** 10.1101/2023.11.10.566641

**Authors:** Guillaume Etter, Suzanne van der Veldt, Coralie-Anne Mosser, Michael E. Hasselmo, Sylvain Williams

## Abstract

The hippocampus is a higher-order brain structure responsible for encoding new episodic memories and predicting future outcomes. In absence of external stimuli, neurons in the hippocampus express sequential activities which have been proposed to support path integration by tracking elapsed time, distance traveled, and other idiothetic variables. On the other hand, with sufficient external sensory inputs, hippocampal neurons can fire with respect to allocentric cues. Previously, these idiothetic codes have been described in conditions where running speed is clamped experimentally. To this day, the exact determinants of idiothetic representations during free navigation remain unclear. Additionally, whether CA1 and CA3 temporal and distance codes are transmitted downstream to the lateral septum has not been established. Here, we develop an unsupervised model trained to compress neural information with minimal loss, and find that we can efficiently decode elapsed time and distance traveled from low-dimensional embeddings of neural activity in freely moving mice. We also developed unbiased information metrics that are minimally sensitive to quantization parameters and enable comparisons across modalities and brain regions. In more than 30,000 CA1 pyramidal neurons, we show that spatiotemporal information is represented as a mixture of idiothetic and allocentric information, the balance of which is dictated by task demand and environmental conditions. In particular, we find that a subset of CA1 pyramidal neurons encode the spatiotemporal distance to rewards. Single cell and population statistics across the hippocampal-septal circuit reveal that idiothetic variables emerge in CA1 and are integrated postsynaptically in the lateral septum. Finally, we implement a computational model trained to replicate real world neural activity, and find that grid cells could provide a plausible input for CA1 representations of time and distance. Altogether, our results suggest that hippocampal CA1 continuously integrates both idiothetic and allocentric signals depending on task demand and available cues, and these high-level representations are effectively transmitted to downstream regions.

## Introduction

The mammalian brain contains a large number of interconnected neurons that not only process incoming sensory information in the present moment, but also perform computations on internally generated, top-down signals that are contingent on past experiences and predictions of the future. Understanding neural activities as a function of this spatiotemporal continuum of experience is an essential question in defining learning mechanisms in biological neural networks. The hippocampus is necessary to encode new experiences [1] and contains neurons that coordinate their activities in stable temporal sequences [2] that represent task relevant information such as spatial locations [3] and elapsed time [4–8] (see Eichenbaum [9] for review). These sequences could in turn provide downstream cortical and subcortical regions with highly associative and context-relevant information [10, 11].

While hippocampal sequences are particularly apparent in linear mazes with stereotyped locomotor patterns [12], they also occur in open environments and track spatial trajectories that can be replayed offline during periods of immobility [13]. In addition to spatial landmarks, the activity of hippocampal neurons can be modulated by salient objects [14], rewards [15] but also abstract, task-relevant features [16].

In hippocampal subfield CA1, these stable internal sequences persist even in absence of external stimuli [17, 18], and are likely the result of developmental priors [19–23]. CA1 sequences could support path integration in absence of cues [24, 25] but anchor to signals from the external world whenever available [14, 26, 27]. Such internal signals could be particularly relevant to bridge epochs of sensory processing, during which information is missing or uncertain. This process can, for instance, be particularly relevant when navigating in darkness [17, 28].

Previously, neurons that tend to fire with respect to internal, idiothetic variables have been related to instrumental measures of time and space [29]. In particular, so-called ‘time cells’ have been defined experimentally by clamping locomotor speed and distance traveled using running treadmills [6, 30]. Importantly, CA1 pyramidal neurons idiothetic representations are not limited to time, but also include distance [6, 31–33] (see McNaughton *et al*. [25] and Mehta [34] for review) and speed [35, 36]. Beyond space and time, the properties of hippocampal neurons have also been defined with respect to internal variables, including network activity [37]. These studies suggest that rather than representing current events, hippocampal neurons organize their activities sequentially to represent experiences on a spatiotemporal continuum.

How idiothetic and allocentric representations contribute to computations that are relevant to downstream regions and memory functions remains unknown. Given that neurons presynaptic to CA1, including CA2 [38] and CA3 [39, 40], could play an integral role in establishing representations of time in CA1, how these representations are then transmitted to downstream postsynaptic regions such as the lateral septum, remains unclear [41].

Here, using calcium imaging of large assemblies of neurons (n *>*30,000) and unsupervised dimensionality reduction of neural signals, we find that complex spatiotemporal representations can be disentangled into distinct allocentric and idiothetic representations in mice freely exploring linear and open environments. At the single neuron level, we develop a non-parametric information theoretic approach and show that CA1 pyramidal neurons always encode a mixture of spatial and temporal information, even outside of tasks involving an explicit temporal component. In addition to tracking past spatiotemporal information, we find that CA1 neurons also encode the time and distance to current goals, and that such representations are enhanced when tracking rewards. Additionally, contextual parameters such as environmental illumination or the presence of auditory tones significantly enhance idiothetic representations. Finally, we find that allocentric representations dominate CA3 activity, while CA1 and lateral septum neurons express idiothetic representations.

## Results

### Hippocampal CA1 neurons encode idiothetic representations during free exploration

To investigate the properties of hippocampal representations in freely moving mice, we transfected large populations of pyramidal neurons in the CA1 subregion with GCaMP6_f_ and recorded calcium activity with open-source miniscopes [42] as animals freely explored linear and open environments (Fig. 1**a**, top). We established the spatial location of neurons in the field of view (Fig. 1**a**, bottom) and for each identified neuron, we extracted the associated calcium transients through time using CNMFe [43] (Fig. 1**b**). We extracted the distance traveled and elapsed time since the onset of each locomotor event, as well as instantaneous speed (1**c**; see Methods). After evaluating correlation coefficients between each allocentric and idiothetic variable pair, we found that time and distance, while correlated at similar traveling speeds, could be disentangled given the diversity of locomotor speeds across runs (Fig. 1**d**; Supplementary Fig. 1).

**Figure 1.**
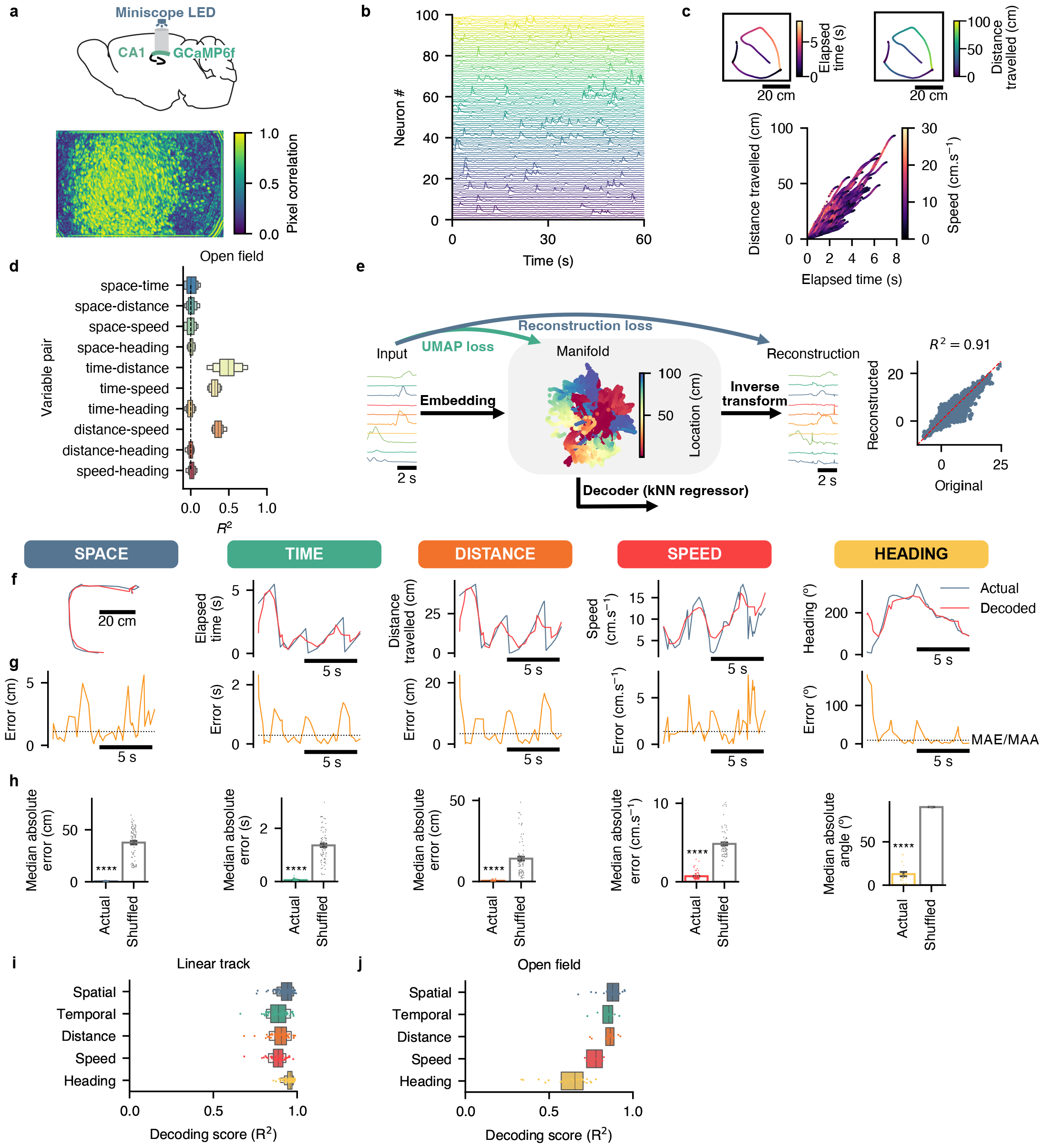
Real-time estimates of CA1 allocentric and idiothetic representations from low-dimensional neural embeddings. **a**, recording configuration (top) and projection of temporally correlated pixels from a calcium imaging video recording (bottom). **b**, calcium transients from a subset of neurons. **c**, example locomotor events in the open field (top) and the relationship between elapsed time and distance traveled for all trajectories for a representative recording session. **d**, low-dimensional embedding method to uncover neural structure over time. Neural recordings are encoded while minimized UMAP loss as well as reconstruction error. Behavioral variables can then be decoded from low-dimensional manifolds using a kNN regressor. **e**, example of spatial signal predicted using a decoder trained on neural embeddings (red line) versus actual labels (blue line). **f**, actual behavioral signals (blue) and predictions from neural embeddings (red). **g**, real-time error between actual and predicted behavioral signals (yellow) as well as median absolute error/angle (MAE/MAA; dotted line). **h**, average error from decoders trained on actual and shuffled behavioral labels. **i**, decoding score (*R*^2^) on the linear track. **j**, same for the open field.

We first focused our analyses of idiothetic representation at the neural population level with the assumption that large ensembles of CA1 neurons contain a certain level of noise and redundancy, and thus their collective representations can be embedded in a number of dimensions that is smaller than the total number of neurons being considered. This unsupervised dimensionality reduction approach presents several advantages over single neuron analyses [44]. In particular, neural manifolds do not require discretization (and thus no bin size parameter), are extracted independently from behavioral signals, and enable comparisons as well as representational alignment between animals. Given these considerations, we designed an embedding method that maximized (1) reconstruction accuracy (i.e. minimal loss when compressing neural data), and (2) interpretability of latent representations (i.e. distinct behavioral states should ideally be represented far apart in latent space). Since time itself was included as a behavioral label, we could not implement contrastive methods that necessitate some form of behavioral labels (including time), such as CEBRA [45]. Using parametric UMAP with an autoencoder loss (pUMAP-AE; [46]), we could obtain low-dimensional embeddings that satisfied the two aforementioned conditions by minimizing both an autoencoder reconstruction loss as well as UMAP loss, while not requiring any behavioral label (Fig. 1**d**; Supplementary Fig. 6; see Methods). We found that we could reliably encode the activity from subsets of n = 128 neurons into 4-dimensional manifolds (see Supplementary Fig. 7 for parameter optimization), suggesting that raw CA1 neural signals contain a mixture of latent representations, correlational redundancy and noise. Importantly, we could reliably decode behavioral signals by training a k nearest neighbor (kNN) regressor on these low-dimensional embeddings. To control for the statistical significance of the decoded behavioral signals, we trained decoders on both actual data as well as shuffled surrogates, where the behavioral label underwent circular permutations (see Methods). Using this method, we were able to decode the location of mice, as well as elapsed time, distance traveled, speed, and heading with sub-second accuracy (Fig. 1**f**). For these variables, we computed the moment-to-moment error between predicted and actual signals, as well as median absolute error (MAE) or median absolute angle (MAA, specifically for heading; Fig. 1**g**). We found that MAE values were higher using shuffled surrogates for position (paired t-test, *t*_83_ = 22.8344, p *<* 0.0001), elapsed time (paired t-test, *t*_83_ = 20.9896, p *<* 0.0001), distance distance traveled (paired t-test, *t*_83_ = 11.6038, p *<* 0.0001), speed (paired t-test, *t*_83_ = 22.0808, p *<* 0.0001), and heading (paired t-test, *t*_18_ = 31.6944, p *<* 0.0001; Fig. 1**h**). We also extracted the decoding score (*R*^2^ values) for each variable in the linear track (Fig. 1**i**) and the open field (Fig. 1**j**). Finally, we computed confusion matrices for each modality across all mice (Supplementary fig. 8).

In addition to a population code for idiothetic representations, we investigated whether CA1 neurons could encode such representations at the single cell level in freely moving conditions. We first binarized the rise periods of calcium transients to extract temporal epochs where neurons are most likely to be active [47] (Fig. 2**a**). Using binarized neural activity and behavioral signals, we computed probabilistic tuning curves for each navigational variable (spatial location, elapsed time and distance traveled since locomotor onset as well as locomotor speed and heading; Fig. 2**b**). We computed each neuron’s information content for a given variable using a metric based on mutual information, corrected for chance, which we term ‘neural information’ (NI; see Methods). NI models random variable associations and corrects values accordingly, is bound between an average of 0 (neural activity has no information about a given variable) and 1 (neural activity perfectly predicts said variable), and has several desirable properties when compared to vanilla mutual information including a reduced dependency on the number of bins used to discretize variables of interest (Supplementary Fig. 2). This latter property allowed us to compare the information content of each neuron across distinct variables irrespective of the number of bins used in the quantization process (see Methods). Additionally, we assessed the significance of neural information using a *χ*^2^ test of independence between neural activity and spatiotemporal variables, and only included data for which p-values were under 0.05 (i.e. significant dependence). We show that the typical distribution of neural information values for distinct variables associated with significant (p ≤ 0.05) is typically above 0.05, and information values below zero (which can occur given that the metric is computed using a null-model with an average of 0) typically do not pass the significance threshold (Supplementary Fig. 3). Additionally, while we found that the onset of locomotion was associated with high correlations between acceleration, time, and distance, neurons encoding time or distance did not systematically encode acceleration (Supplementary Fig. 4).

**Figure 2.**
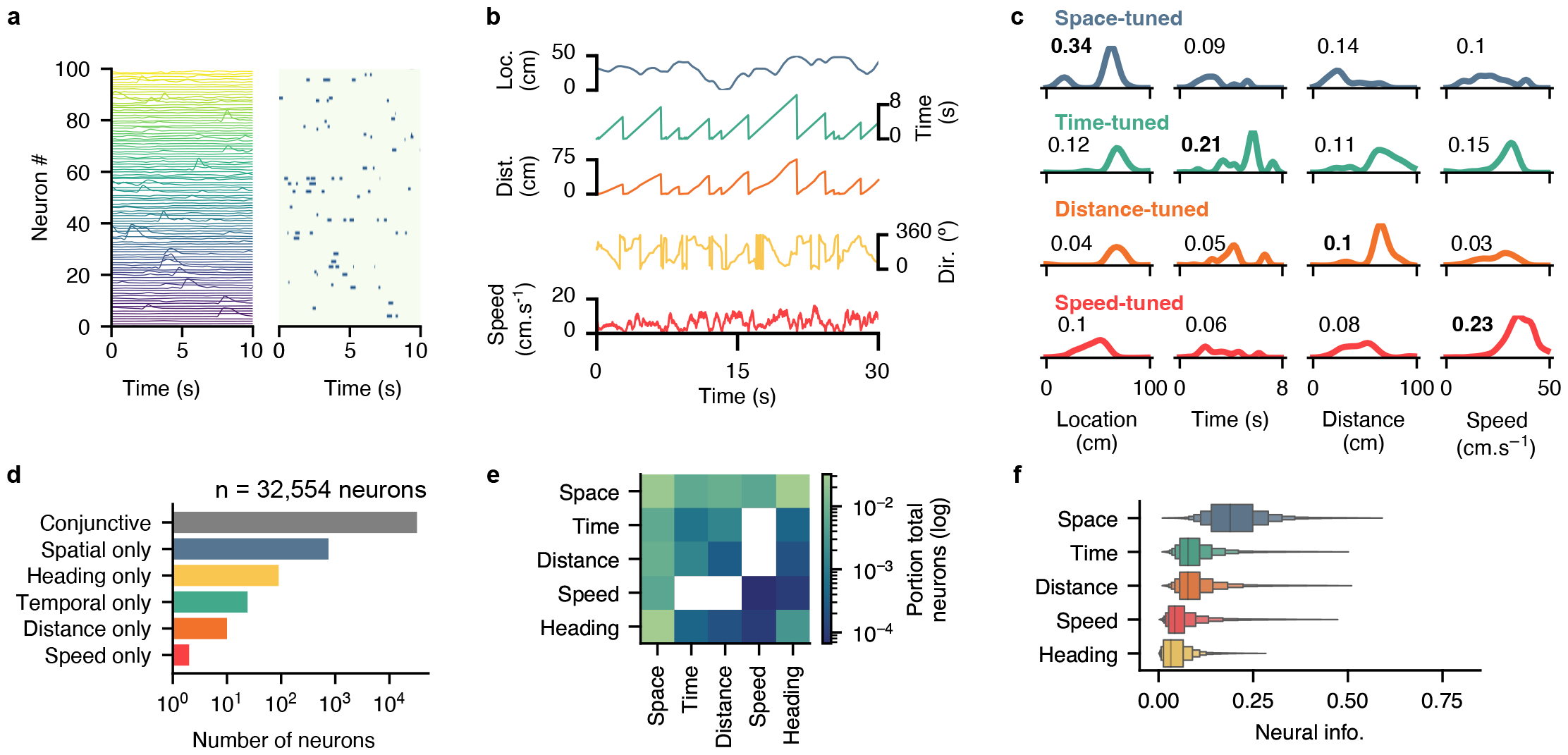
CA1 single cell representations of idiothetic and allocentric features in freely behaving conditions. **a**, calcium transients and corresponding binarized activity for an example recording epoch. **b**, location (blue), time elapsed (green), distance traveled (orange), heading (yellow), and speed (red) for an example recording epoch. **c**, representative example tuning curves to space, time, distance, and speed. Each row is an example cell most prominently tuned to space (blue), time (green), distance (orange) or speed (red). Top left numbers indicate the peak activity likelihood for each neuron at a given discretized state. **d**, number of neurons tuned to one variable only, with the exception of conjunctive neurons (gray) that are tuned to at least one variable (or more). **e**, portion of neurons tuned to two variables conjunctively. The diagonal represents the portion of neurons tuned to only one variable. **f**, neural information for each variable.

In addition to space, we found that a subset of CA1 neurons could encode idiothetic variables such as time, distance, or speed in freely moving conditions (Fig. 2**c**). The vast majority of neurons conjunctively encoded more than one variable (Fig. 2**d**), with many encoding a mixture of features conjunctively (Fig. 2**e**). When considering the overall information content of CA1 neurons, we found that space was most prominently encoded (0.1975 *±* 0.0005 NI), followed by distance (0.0825 *±* 0.0003 NI), time (0.081 *±* 0.0003 NI), speed (0.0404 *±* 0.0002 NI), and finally heading direction (0.0341 *±* 0.0002 NI; Fig. 2**f**). Interestingly, we found that neurons tended to encode space more prominently in open (0.2626 *±* 0.0007) versus linear (0.1731 *±* 0.0005) environments, and heading was encoded more prominently in open (0.0746 *±* 0.0004) versus linear (0.019 *±* 0.0002) environments (2-ANOVA, *F*_4_ = 7018.7970 for the interaction between modality and maze type; p *<* 0.0001; Supplementary Fig. 5,**b**). Altogether, these findings confirm that individual CA1 neurons can represent idiothetic information, including time, distance, and speed, during free exploration.

### Darkness and auditory cues enhance CA1 idiothetic representations

We next investigated whether behavioral constraints could modulate the predominance of idiothetic versus allocentric representations in hippocampal CA1 neurons. Previous studies suggested that the tuning of CA1 neurons to time and distance could emerge in darkness [17, 28]. Thus, we first compared the information content of CA1 neurons of mice exploring a linear track in either lit or dark conditions. We found that environmental illumination had a significant influence on spatiotemporal information on the linear track (2-ANOVA, *F*_4_ = 45.46, p < 0.0001 for the interaction between modality and illumination condition; Fig. 3**a**), with CA1 neurons displaying stronger tuning to idiothetic signals compared to lit environments. We found that CA1 neurons contained more information about time elapsed when running in darkness (0.0965 *±* 0.0026) compared to lit environments (0.0849 *±* 0.0003; pairwise t-test, *t*_749.24_ = 5.9118, p *<* 0.0001). These neurons also encoded distance traveled better in darkness (0.1005 *±* 0.003) compared to lit environments (0.086 *±* 0.0003; pairwise t-test, *t*_751.94_ = 6.9152, p *<* 0.0001). Similarly, speed was also better encoded in darkness (0.0488 *±* 0.0018) compared to lit environments (0.0432 *±* 0.0002; pairwise t-test, *t*_567.27_ = 4.2914, p *<* 0.0001). On the other hand, allocentric location was poorly encoded in darkness (0.1969 *±* 0.0034) compared to lit environments (0.2037 *±* 0.0004; pairwise t-test, *t*_868.38_ = 1.9957; p = 0.0463) although this statistical significance was associated with small statistical power (Hedges *g* = 0.0820). Finally, CA1 neurons contained more information about heading direction in well-lit (0.0475 *±* 0.0003) compared to dark environments (0.0276 *±* 0.0011; pairwise t-test, *t*_555.55_ = 17.2017, p *<* 0.0001; Fig. 3**b**). A confusion matrix for heading direction decoded from low-dimensional manifolds also revealed that decoding errors increased in darkness conditions (Fig. 3**c**).

**Figure 3.**
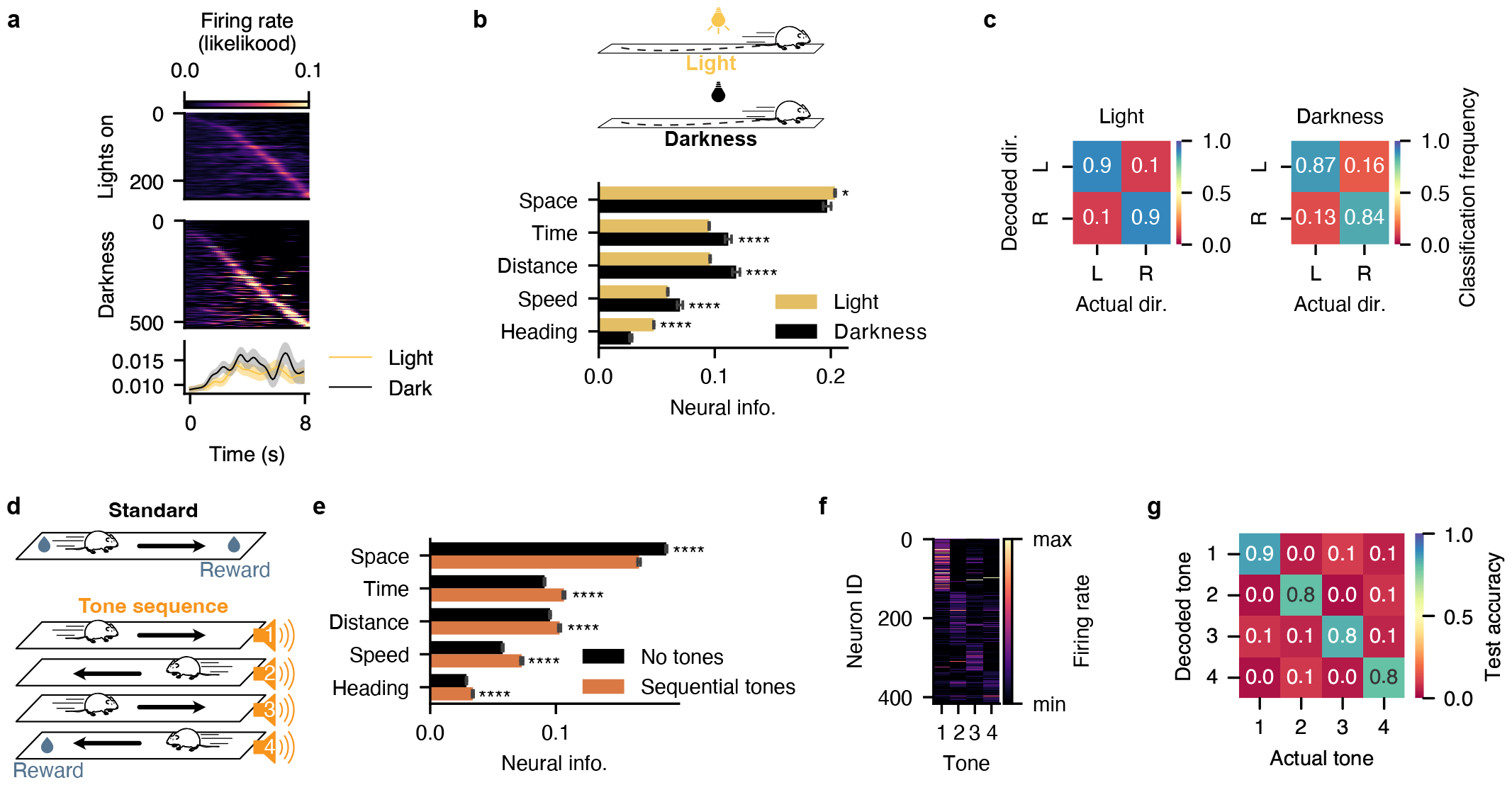
CA1 idiothetic representations are modulated by contextual conditions. **a**, tuning curves for neurons encoding time significantly, sorted by preferred firing time after the onset of locomotion, for both light and darkness conditions. Probabilistic firing rate (likelihood) as a function of elapsed time in light (yellow) and darkness (black). Error bands represent SEM. **b**, neural information for spatiotemporal signals in both while mice explored a linear track in either the presence of ambient light or in darkness. **c**, confusion matrices for the decoded linear track running direction in light (left) and darkness (right). Red, low classification frequency; dark blue, high classification frequency. **d**, mice were trained on either a standard linear track, or one that was augmented with a sequence of tones (4 auditory contexts) that lead to the delivery of a reward. **e**, neural information for spatiotemporal signals on the linear track, in presence or absence of a tone sequence. **f**, tuning curves for a subset of neurons that are significantly tuned to a tone in the tone sequence. **g**, confusion matrix for the decoding accuracy of tones.

In a previous study, we assessed spatiotemporal tuning in a linear maze where sequential tones were presented in order to extend the length of behavioral sequences leading to reward consumption [33] (3**d**). Here, we find that adding a tone sequence to the linear track task changes significantly the spatiotemporal representations of CA1 neurons (2-ANOVA, *F*_4_ = 265.6141, p *<* 0.0001 for the interaction between modality and presence of tones). In particular, temporal information was lower in a classical linear track (0.0787 *±* 0.0022) compared to the one augmented with sequential tones (0.1064 *±* 0.0009; pairwise t-test, *t*_7007.22_ = 15.4236, p *<* 0.0001). Similarly, distance information was lower in the classical linear track (0.0824 *±* 0.002) compared to the one augmented with sequential tones (0.1029 *±* 0.001; pairwise t-test, *t*_6809.31_ = 7.0104, p *<* 0.0001). Speed information was lower in the classical linear track (0.0533 *±* 0.0023) compared to the one augmented with sequential tones (0.0729 *±* 0.001; pairwise t-test, *t*_4936.14_ = 13.5966, p *<* 0.0001). Interestingly, spatial representations contained more information in the regular linear track (0.1714 *±* 0.0037) compared to the one augmented with sequential tones (0.1662 *±* 0.0011; pairwise t-test, *t*_7943.88_ = 17.1003, p *<* 0.0001; Fig. 3**e**). Additionally, we found that CA1 neurons could encode specific tones on the linear track augmented with sequential tones (3**f**). We found that a subset of neurons could encode a specific tone, and that we could successfully decode the exact tone in the sequence from low-dimensional neural embeddings (3**g**).

### Current goals modulate prospective representations of time and distance

Beyond environmental conditions, we next evaluated whether current goals could modulate idiothetic representations. In particular, we made a distinction between time and distance ‘to’ (prospective), versus ‘from’ (retrospective) reward locations. We first compared a standard linear track where rewards are delivered at either end of the track, to a linear track where a tone is triggered when the mouse reaches one side and a reward is delivered only on the opposite side (Fig. 4**a**). Importantly, triggering this tone is required to obtain a reward on the opposite side of the linear track. While mice performed this task, we assessed both the time and distance since the onset of each trajectory event (retrospective) as well as the time and distance until the end of the ongoing trajectory event (prospective; Fig. 4**b**). Using these signals and binarized neural signals, we computed tuning curves to both retrospective as well as prospective time and distance, which revealed that a subset of neurons could encode specifically each of these signals (Fig. 4**c**).

**Figure 4.**
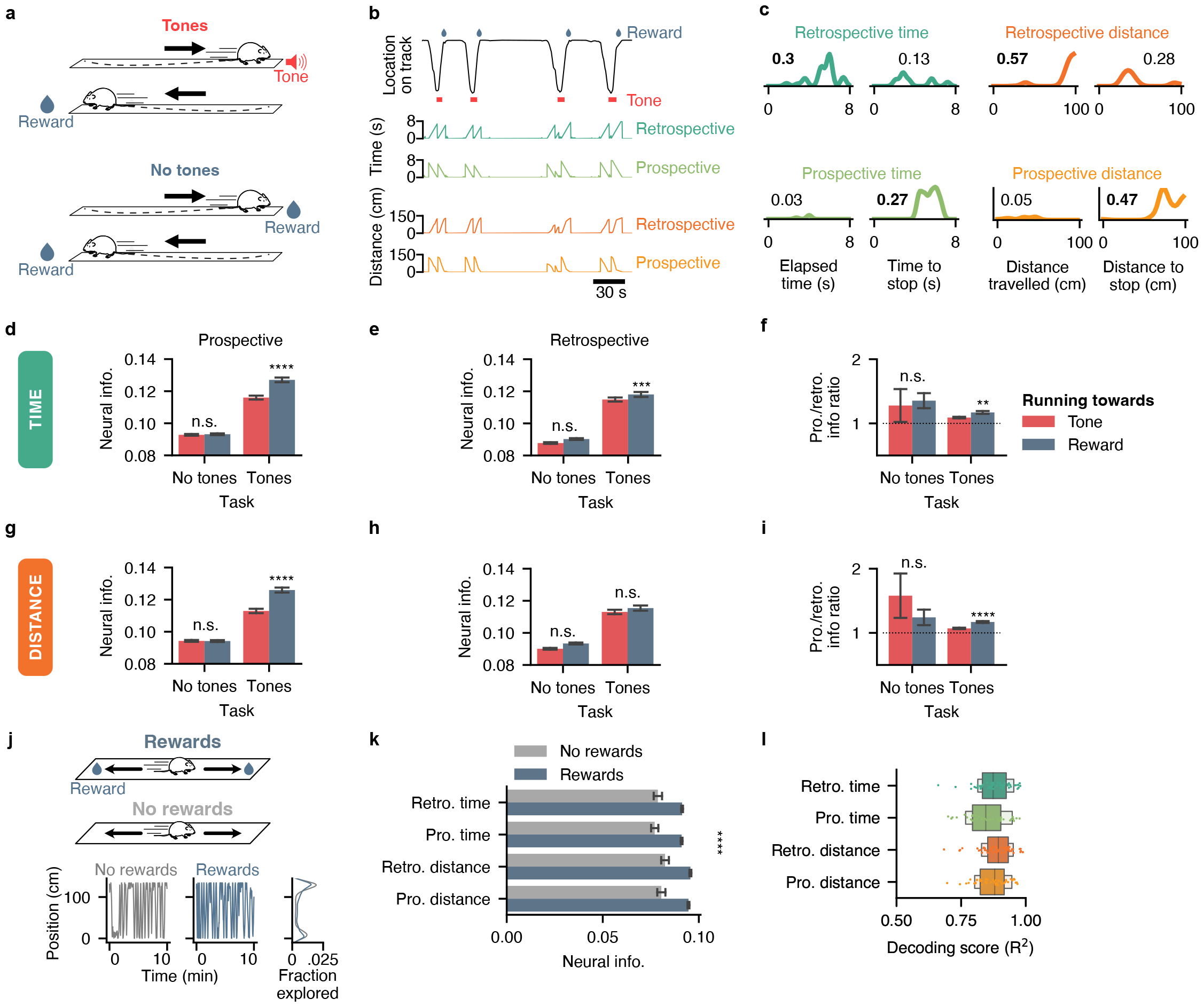
Current goals dictate the prevalence of CA1 prospective representations. **a**, mice were trained to either collect rewards on either end of a linear track, or to alternate between triggering a tone on one end and collecting a reward on the opposite end. **b**, for each running trajectory, both prospective and retrospective time and distance were registered along with neural signals. **c**, example tuning curves for neurons encoding prospective or retrospective time or distance specifically. **d**, average neural information for prospective time in a standard linear track (‘No tones’ condition) versus a linear track where one reward is replaced with a tone. Running trajectories towards rewards (blue) or towards tones (red) are distinguished. **e**, same for retrospective time. **f**, ratio of prospective over retrospective time. **g**, average neural information for prospective distance in either task. **h**, same for retrospective distance. **i**, ratio of prospective over retrospective distance. **j**, mice were trained to run to either end of a linear track in presence of absence of rewards to collect (top). Bottom, linear track exploration for one example mouse in a session that included rewards (blue) and one that did not include rewards (gray), as well as the corresponding histogram distribution of explored locations for both sessions. **k**, neural information for each modality in presence (blue) or absence (gray) of rewards to collect. **l**, decoding error for prospective and retrospective time and distance using low-dimensional neural embeddings.

To investigate how task demands modulated the prospective or retrospective nature of these idiothetic signals, we quantified neural information for each CA1 neuron while distinguishing trajectories by goals: trajectories towards rewards (and away from the zone that triggers tones) versus trajectories away from rewards (and towards the zone that triggers tones). Importantly, we found an interaction between the running direction and the task used (2-ANOVA, *F*_1_ = 36.9905, p < 0.0001 for the interaction between running direction and task): prospective temporal information was higher when mice ran towards rewards (0.127 *±* 0.0014) compared to when mice ran towards the tone-triggering area (0.116 *±* 0.0012; pairwise t-test, *t*_4814.35_ = 5.8205, p *<* 0.0001). As a control, running direction had no influence on prospective temporal information in a regular linear track that did not contain tones (pairwise t-test, *t*_18701.50_ = 0.4810, p = 0.6305; Fig. 4**d**). On the other hand, retrospective temporal information (i.e. time since onset of locomotion) was also associated with a significant interaction between running direction and task type (2-ANOVA, *F*_1_ = 9.1293, p = 0.0025) and retrospective temporal information was also higher when mice ran towards rewards (pairwise t-test, *t*_4728.75_ = 3.4756, p = 0.0004) while running direction had no influence on retrospective temporal information in a regular linear track that did not contain tones (pairwise t-test, *t*_18384.06_ = 1.8410, p = 0.0656; Fig. 4**e**). To summarize this analysis, we also compute the ratio of prospective over retrospective information and found that this ratio only increased significantly (from 1.09 to 1.17) in the task that involved alternating between a tone and a reward (pairwise t-test, *t*_3250.30_ = 3.1582, p = 0.0016; 4**f**).

Additionally, we investigated whether distance signals displayed similar properties in CA1 neurons. As for temporal information, we found a significant interaction between the running direction and type of task used (2-ANOVA, *F*_1_ = 47.5735, p *<* 0.0001), and CA1 neurons encoded more prospective distance information when running towards rewards (0.126 *±* 0.0015) when compared to running towards the tone-triggering area (0.1129 *±* 0.0014; pairwise t-test, *t*_4795.5532_ = 6.3466, p *<* 0.0001; 4**g**). However, unlike for retrospective temporal information, there was no interaction between running direction and the task type for retrospective distance information (2-ANOVA, *F*_1_ = 0.7366, p = 0.3908). For both the classical linear track and the one augmented with a tone, retrospective distance information was higher when running towards rewards (2-ANOVA, *F*_1_ = 13.3601, p = 0.0003; 4**h**). When computing the ratio of prospective over retrospective distance information, we found that this ratio only increased significantly (from 1.07 to 1.17) in the task that involved alternating between a tone and a reward (pairwise t-test, *t*_4107.77_ = 5.5872, p *<* 0.0001; 4**i**).

Since CA1 neurons are known to fire with respect to reward locations [15], we also evaluated the relationship between spatiotemporal information and the presence or absence of rewards on the linear track. Since mice already learned the standard linear track task by the time we removed rewards, we found that linear track exploration was essentially comparable to rewarded exploration (Fig. 4**j**). The presence of rewards increased the overall tuning of CA1 neurons to idiothetic signals (2-ANOVA, *F*_1_ = 75.6663, p = 0.0001 for the main effect of reward presence; 4**k**). Importantly, we were also able to decode both prospective and retrospective idiothetic representations from low-dimensional neural manifolds (Fig. 4**l**)

### Computational mechanisms underlying idiothetic representations in freely moving conditions

As CA1 neurons receive both place cell inputs from CA3 as well as grid cell inputs from the medial entorhinal cortex, we investigated what presynaptic inputs could support the emergence of idiothetic representations in CA1. To this end, we simulated place and grid cells using real trajectories from mice exploring linear and open environments. The respective activity rate from both cell types was then used to train an artificial model neuron to predict real activity of each individual neuron associated with the exploration trajectory. Specifically, synaptic inputs are integrated in a linear manner [48, 49]. To evaluate the capacity of our model to leverage grid or place cell inputs, we computed F_1_ scores for each neuron using either a place or grid cell input model (Fig. 5**a**). To evaluate how distinct input representations could best explain idiothetic versus allocentric tuning, we compared F_1_ scores using place or grid cells inputs to top-50 neurons for allocentric and idiothetic signals. On the one hand, we found that a grid cells input model could significantly better predict the activity of time (pairwise Tukey, F_49_ = 12.3241, p = 0.0003) and distance (F_49_ = 17.2270; p *<* 0.0001) neurons compared to input place cells. On the other hand, the activity of neurons most tuned to heading direction was best explained by a place cell input model (F_49_ = 10.3301, p = 0.0007; Fig. 5**b**). We also computed a grid-over-place fit ratio (where 1 and -1 corresponds to neural activity being exclusively explained by input grid or place cells, respectively). We find that both time and distance are significantly better explained by input grid cells (1-ANOVA, F_49_ = 23.1524, p *<* 0.0001), whereas place cell activity could be explained equally by either grid or place cell models (Fig. 5**c**). Altogether, this suggests that idiothetic representations in freely moving mice could arise from the integration of presynaptic grid cell inputs.

**Figure 5.**
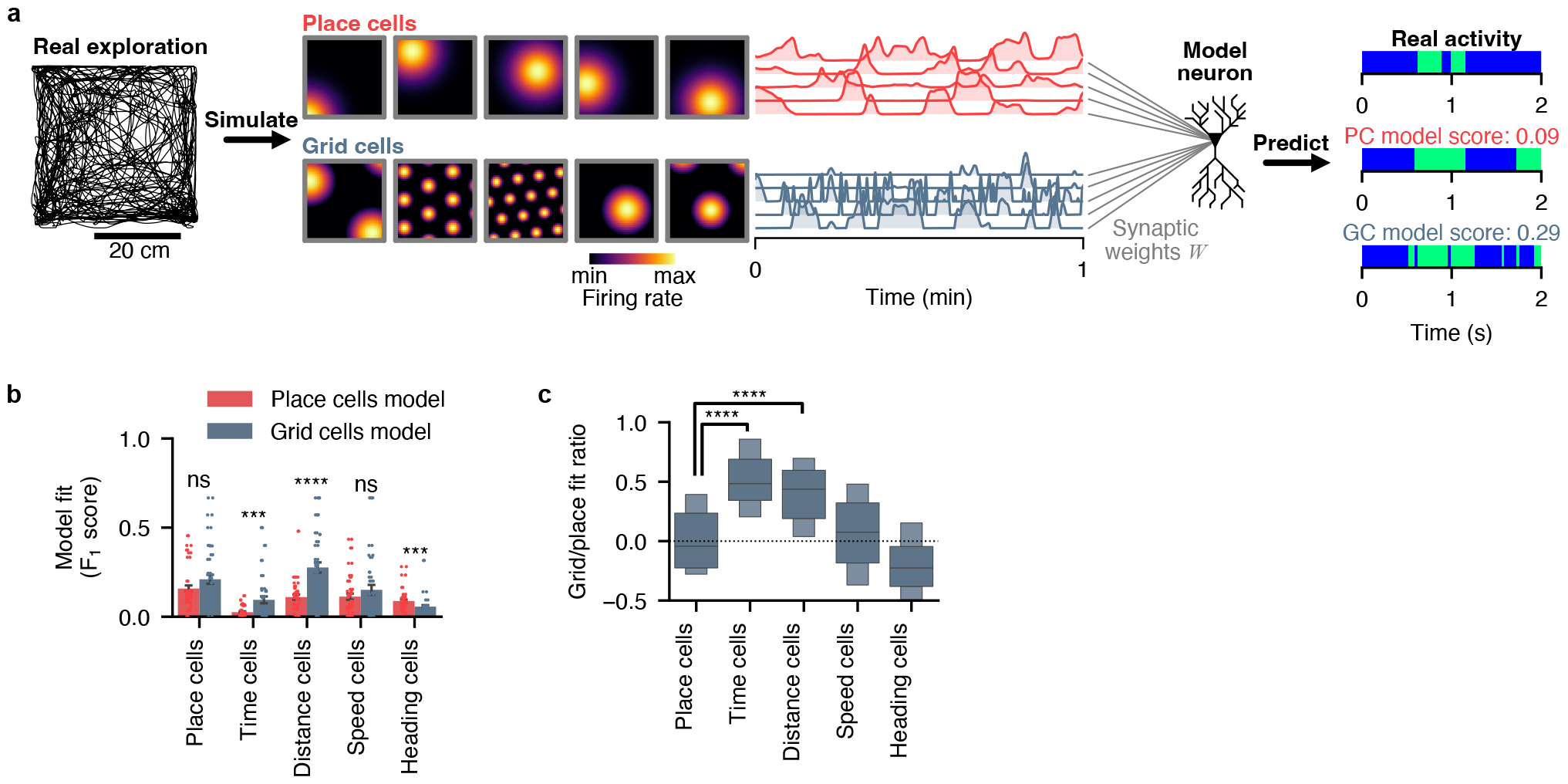
Computational mechanisms underlying idiothetic representations in freely moving conditions. **a**, approach for estimating the relative contribution of place and grid cells to idiothetic signals. Place and grid cell rate maps are generated from a real exploration trajectory. The respective activity rate from both cell types are then used to train a logistic regressor to predict the real activity of each individual neuron. Finally, the F_1_ score is computed for each neuron using either a place or grid cell input model. **b**, F_1_ scores for place (red) or grid (blue) cell inputs used to predict the top-50 neurons exclusively tuned to space, time, distance, speed, or heading direction. **c**, grid-over-place fit ratio for each cell type.

### Integration of spatiotemporal representations from CA3 and transmission to the lateral septum

It was previously proposed that CA1 spatial and temporal representations could be inherited from CA3 and CA2 respectively [38, 50]. As the lateral septum is the main sub-cortical output from the hippocampus (receiving both inputs from CA1 and CA3) [51], we next sought to relate the properties of spatiotemporal codes extracted in CA1 neurons, to those of presynaptic CA3 and postsynpatic lateral septum neurons. For each of these three brain regions, we performed calcium imaging (Fig. 6**a**), extracted spatial footprints of neurons expressing GCaMP6f (Fig. 6**b**,**c**) as well as corresponding calcium transients (Fig. 6**d**). In turn, for each brain region, we extracted low-dimensional embeddings (using an input of *n* = 32 neurons, mapped into *D* = 4 latent dimensions for every mouse, to account for the lower maximum sampling in CA3 mice; Fig. 6**e**) which we later use to compare representations across animals recorded in distinct regions for both the linear track and open-field, in normal conditions (well-lit environments with no tones). Notably, there was a significant interaction between brain regions and the modalities encoded (2-ANOVA, *F*_8_ = 285.7025, p *<* 0.0001 for the interaction between regions and modalities): CA3 neurons contained significantly more information about space (0.2617 *±* 0.0027) compared to CA1 (0.1923 *±* 0.0004; pairwise t-test, *t*_1005.1494_ = 25.9814, p *<* 0.0001) or lateral septum neurons (0.1822 *±* 0.0012; pairwise t-test, *t*_1389.3390_ = 27.2796, p *<* 0.0001). In contrast, neurons in CA1 displayed better representations of time (0.0855 *±* 0.0003) compared to CA3 (0.0524 *±* 0.0012; pairwise t-test, *t*_1065.0931_ = 26.5503, p *<* 0.0001) and lateral septum neurons (0.0603 *±* 0.0005; pairwise t-test, *t*_11172.0104_ = 45.7872, p *<* 0.0001; Fig. 6**f**). Interestingly, the stability of low-dimensional neural embeddings (within-session train-test manifold reliability) was significantly lower in CA1 (∼0.3), when compared to either CA3 or lateral septum recordings (∼0.6; 1-ANOVA, *F*_2_ = 139.8172, p *<* 0.0001, *η*^2^ = 0.523, n = 57 independent mice; Fig. 6**g**).

**Figure 6.**
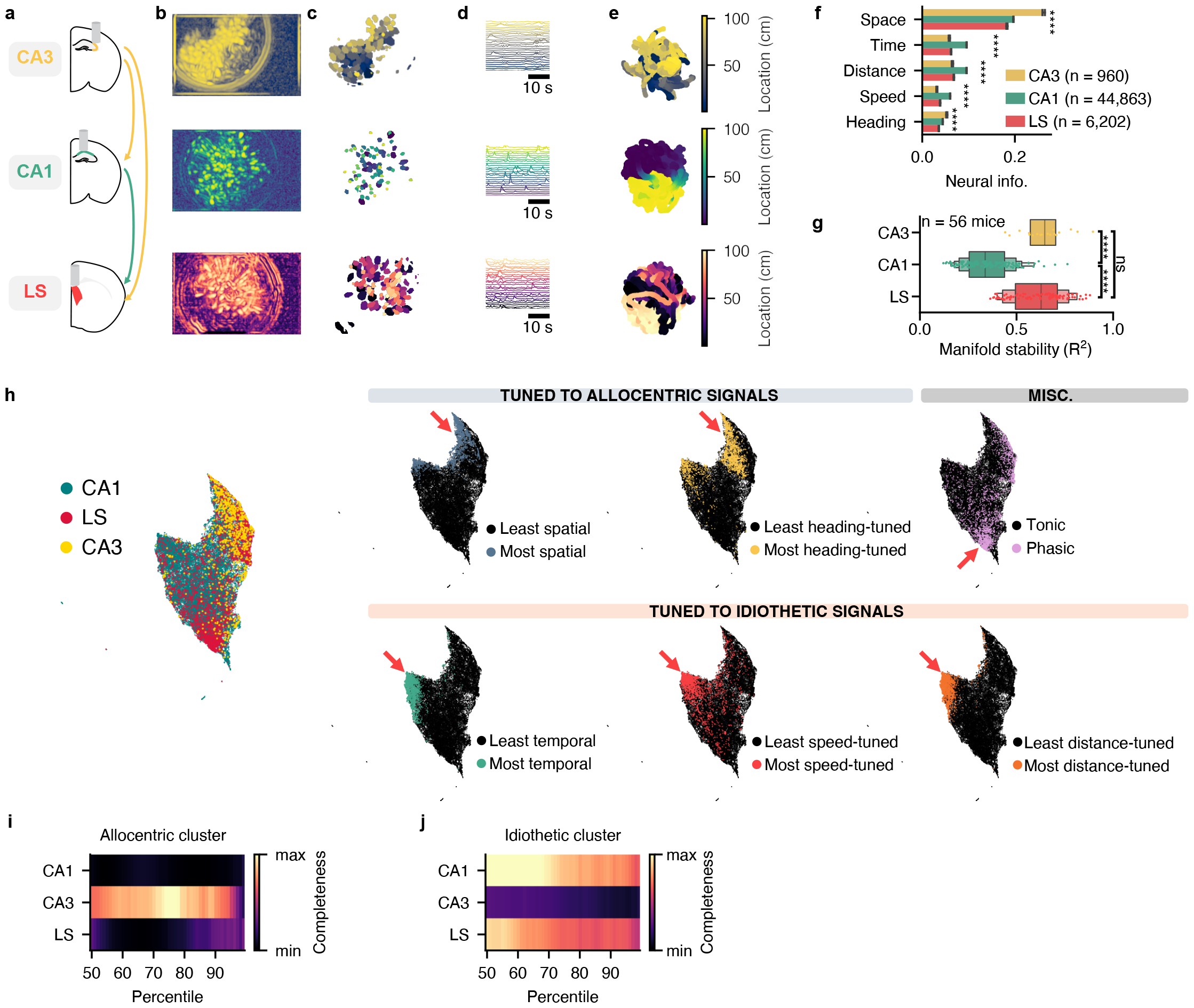
Integration of spatiotemporal representations from CA1 and CA3 in the lateral septum. **a**, recording configuration in CA3 (yellow), CA1 (green), and the lateral septum (red). **b**, projections of temporally correlated pixels for example recordings in each region. **c**, corresponding spatial footprints extracted in each region. **d**, subset of calcium transients extracted in each region. **e**, corresponding low-dimensional manifolds extracted in each region. Displaying only two out of four dimensions for clarity. **f**, neural information for spatiotemporal variables in each region. **g**, within-session manifold stability (*R*^2^) for mice recorded in CA3, CA1, or the lateral septum. **h**, unsupervised, low-dimensional projection of neurons from all regions based on spatiotemporal and firing properties. Each dot represents a neuron, and color codes are used to display distinct labels. **i**, completeness score between region labels and n^*th*^ percentile of neurons encoding allocentric variables the most. **j**, same but for idiothetic signals.

We next sought whether each brain region could be clustered using an unsupervised machine learning method, by embedding each neuron using a set of basic properties (tuning to space, time, distance, speed, heading, activity rate, transition probabilities; see Methods). After projecting each neuron on a low-dimensional manifold based on these properties using vanilla UMAP, we applied a color code for each region and for neurons corresponding to the 95^*th*^ percentile of each property value (Fig. 6**h**). We found two major cell clusters corresponding to neurons encoding either allocentric or idiothetic variables predominantly. On the one hand, CA3 neurons were associated with neurons that best encode allocentric variables (Fig. 6**i**). On the other hand, speed was most associated with the neurons in the lateral septum (Fig. 6**j**). Altogether, these results suggests that CA3 neurons encode allocentric representations predominantly, while the lateral septum inherits a mixture of allocentric and idiothetic representations from CA1 and CA3.

## Discussion

To this day, how neurons in the hippocampal-septal circuit integrate allocentric and idiothetic signals remains unclear. Here, we found that we could disentangle spatiotemporal codes in freely moving animals exploring linear as well as open environments and that both allocentric and idiothetic representations could be decoded from low-dimensional neural embeddings. In particular, we find that CA1 pyramidal neurons can track elapsed time and distance traveled during independent bouts of locomotor activity. We also find that contextual conditions, such as the absence of environmental illumination and task cues, can enhance idiothetic representations relative to allocentric ones. Additionally, CA1 neurons can encode prospective idiothetic representations that are associated with runs towards rewards. We find that in the hippocampal-septal system, CA1 and CA3 predominantly encode idiothetic and allocentric representations, respectively. While both streams of information are integrated downstream in the lateral septum, representational stability is highest in CA3 and the lateral septum. Finally, training a model on real-work exploration and neural activity, as well as simulated place and grid cell inputs, we find that grid cells could constitute a plausible source of activity for the generation of prospective and retrospective idiothetic representations.

### Temporal representations can be expressed outside of memory tasks

To assess temporal representations, previous studies have typically used a running treadmill [5, 6, 30, 52] or virtual reality [31, 32, 53] to clamp the spatial location of animals and assess the tuning of individual neurons to discrete temporal windows. Additionally, generalized linear models applied to such data have revealed that hippocampal activity cannot be entirely explained by spatial variables, and elapsed time significantly contributes to hippocampal activity [6]. Here, using an adjusted information theoretic metric, we show that it is possible to extract the relative information content of individual neurons with respect to allocentric and idiothetic variables. We also establish how time, distance, and speed co-vary in linear and open environments, and establish the relative population statistics for specific neurons (that encode one variable but not others) and conjunctive neurons (that encode more than one variable), which is compatible with previous observations that CA1 representations are largely multiplexed [53]. From a computational standpoint, mixed selectivity is a desirable property that supports robust representational encoding since it enables a high degree of separability for represented modalities [54] (see Fusi *et al*. [55] for review).

### Idiothetic representations as primordial features of the hippocampal predictive map

While early interpretations of hippocampal representations have been related to a ‘cognitive map’ that registers the geometry of surroundings ([56]), more recent interpretations propose that the hippocampus learns a predictive map ([57]). This model, termed ‘successor representation’, proposes that neurons in the hippocampus represent the value of future states instead of ‘pure’ spatial locations. This is a reasonable model as place fields tend to not represent perfectly the external world and instead display asymmetry over the course of learning ([58]) and sensitivity to rewards ([59]). CA3 neurons were also found to specifically express prospective representations on short, compressed timescales [60]. Here, we found that in addition to previously described retrospective idiothetic representations, CA1 neurons can encode prospective idiothetic signals (i.e. time and distance towards future goals). Interestingly, the ratio of prospective to retrospective information was systematically higher when running towards rewards, compared to running towards auditory cues. While the hippocampus has previously been shown to track past, present, and future locations on a sub-second scale [61], our results indicate that CA1 neurons could represent past and future states using idiothetic temporal and distance information. More specifically, rather than tracking a future location relative to the current one, CA1 neurons could specifically track the time or distance to a future goal. These predictive representations could act as representational priors, particularly when spatial sensory evidence is lacking. As described previously [28], we found that idiothetic representations were enhanced in darkness, as well as in behavioral tasks including non-spatial, auditory cues. In the latter case, we observed that some CA1 neurons tune their activities to specific contextual tones, similar to previous descriptions [62].

Previously, it was proposed that the temporal organization of hippocampal activities depends on medial entorhinal cortex inputs [63, 64]. Our computational model suggests that idiothetic representations could be inherited from presynaptic entorhinal grid cells, which is in contrast to hippocampal place cells that have been proposed to rely on external sensory inputs, more so than grid or boundary vector cells [65]. This idea is supported by the observation that grid cells display activity that is inherently periodic, and already contain representations of time [66], and speed [67, 68]. One notable limitation of our model is the omission of theta oscillations in the generation of grid cells. It should be noted that oscillatory interference models used to generate grid fields add pertinent properties to the model, such as context-dependent grid firing upon phase resetting, and could play an important role in supporting temporal representations during navigation [69]. Moreover, our proposal also has to be balanced by the observation that time cells specifically expressed during immobility [70] can subsist after medial entorhinal cortex lesions [71], suggesting that these temporal representations could depend on other regions or emerge through local interactions.

### Generalization of hippocampal population representations across the hippocampal-septal circuit

While our analysis of tuning curves shows that a large number of CA1 neurons fire with respect to prospective or retrospective idiothetic signals, analyses at the single neuron level present some limitations. In particular, individual tuning curves need to be discretized and cannot be generalized across animals, as tuning curves computed in one animal do not have predictive values for other animals. To solve this problem, previous studies have introduced methods to extract low-dimensional embeddings of neural data [16, 37, 72] assuming a certain amount of redundancy and noise in neural data. However, these techniques do not explicitly minimize compression loss in their objective function. Other approaches relying on contrastive learning objectives provide reliable embeddings while minimizing reconstruction loss, and can use time vectors as training labels in their least supervised form [45]. Since time itself was a hypothetical variable encoded by neural ensembles in this study, we chose to implement a truly unsupervised objective function instead. Here, by leveraging parametric UMAP with an autoencoder loss, we propose a simple embedding method that maximizes both latent interpretability (UMAP objective) and minimizes loss in neural data (reconstruction objective) without requiring training labels. We find that manifolds extracted with this method outperform vanilla UMAP, and enable precise decoding of spatiotemporal variables. Finally, we explore how spatiotemporal representations in CA1 could be inherited from presynaptic neurons in CA3, and transmitted to postsynaptic neurons in the lateral septum. While we find that CA3 also encodes time (as previously shown [73]) and distance, CA3 neurons exhibit the strongest levels of spatial and heading information. On the other hand, while it was previously described that the lateral septum could express a spatial code dependent [74] and independent [75, 76] on the phase of hippocampal theta oscillations, we find that the lateral septum can represent time, distance, as well as speed, which could be relevant in evaluating future decisions on short timescales [77]. Previously, it was noted that CA3 neurons exhibited enhanced representational stability when compared to CA1 [50]. Here we found using low dimensional neural embeddings that while the lateral septum receives inputs from both CA1 and CA3, its representational stability is similar to that of CA3, and drastically higher than CA1, which confirms single cell properties previously described [75]. We attribute this lower CA1 manifold stability to representational drift, which has recently been well-documented in CA1 pyramidal neurons [78–80].

Altogether, our results indicate that CA1 pyramidal neurons encode idiothetic signals including time, distance, and speed. The prevalence of these signals over allocentric representations of space and direction is directly dependent on environmental conditions and task demands. Representing future goals such as rewards could play a major role in navigation, learning, and memory.

## Methods

### Animals

All procedures were approved by the McGill University Animal Care Committee and the Canadian Council on Animal Care (protocol 2015-7650). A total of *n* = 56 male (*n* = 29) and female (*n* = 28) 8–16 weeks old, B6;129P2 PV-Cre mice (Jackson Laboratory, RRID:IMSR_JAX:017320), VGAT-IRES-Cre (Slc32a1tm2(cre)Lowl/J, Jackson Laboratory, JAX:016962), C57bl/6 (Jackson Laboratory, JAX:016962) and Grik4-Cre (C57BL/ 6-Tg(Grik4-cre)G32-4Stl/J, Jackon Laboratory, JAX :006474) were used in this study. Mice were housed individually on a 12-h light/dark cycle at 22 °C and 40% humidity with food and water ad libitum. All experiments were carried out during the light portion of the light/dark cycle.

### Adeno-associated viral vectors

Adeno-associated virus AAV5.CamKII.GCaMP6f.WPRE.SV40 (Addgene # 100834, obtained from the University of Pennsylvania Vector Core) or AAV2/9.Syn.Flex.GCaMP6f.WPRE.SV40,# CS0641, obtained from the University of Pennsylvania Vector Core) was used in all calcium imaging experiments.

### Surgical procedures

Mice were anesthetized with isoflurane (5% induction, 0.5–1% maintenance) and placed in a stereotaxic frame (David Kopf Instruments). Body temperature was maintained with a heating pad, eyes were hydrated with gel (Optixcare), and Carprofen (10 ml/kg) was administered subcutaneously. The skull was completely cleared of all connective tissue and small craniotomies were performed using a dental drill for subsequent injection or implant.

### Viral injections

All viral injections were performed using a glass pipette connected to a Nanoject III (Drummond) injector. We injected the AAV5.CamKII.GCaMP6f virus (200 nL at 1 nL.s^−1^) in hippocampal CA1 using the following coordinates: anteroposterior (AP) -1.86 mm from bregma, mediolateral (ML) 1.5 mm, dorsoventral (DV) 1.5 mm [81]. For lateral septum targeted injections, we injected AAV2/9.Flex.GCaMP6f using the following coordinates: Anterior LS, AP: 0.86 mm; ML: 0.35 mm; DV: -3.0 mm; Intermediate LS, AP: 0.38 mm; ML: 0.35 mm; DV: -2.7 mm; Posterior LS, AP: 0.10 mm; ML: 0.35 mm; DV: - 2.5 mm, with DV *±* 0.2 to adjust for more dorsal or ventral placement of injection. LS injections were done in VGAT-IRES-Cre transgenic mice. For dorsal CA3 targeted injections, C57Bl/6J mice were injected with AAV5.CamKII.GCaMP6f at AP: -2.1 mm; ML: 2.3 mm; DV: -2.2 mm. Three of the included CA3 animals were Grik4-Cre transgenic mice injected with AAV2/9.Flex.GCaMP6f, restricting GCaMP6f expression to the pyramidal cells of CA3 specifically.

### Implants for calcium imaging

Two to four weeks post GCaMP6f injection, a 0.5 mm diameter gradient refractive index (GRIN, Inscopix, Palo Alto, California, United States of America) lens was implanted above LS, dorsal CA1 or CA3, or a 1.8 mm GRIN (Edmund Optics, Barrington, New Jersey, United States of America) lens was implanted above dorsal CA1. For 1.8 mm CA1 implants, mice were anesthetized with isofluorane and the skull was cleared. A ∼2 mm diameter craniotomy was performed in the skull above the injection site. An anchor screw was placed on the posterior plate above the cerebellum. The dura was removed, and the portion of the cortex above the injection site was aspirated using a vacuum pump until the corpus callosum was visible. These fiber bundles were then gently aspirated without applying pressure on the underlying hippocampus, and a 1.8 mm diameter gradient index (GRIN; Edmund Optics) lens was lowered at the following coordinates: AP -1.86 mm from bregma, ML 1.5 mm, DV 1.2 mm. The GRIN lens was permanently attached to the skull using C&B-Metabond (Patterson Dental), and Kwik-Sil (World Precision Instruments) silicone adhesive was placed on the GRIN lens to protect it. Four weeks later, the silicone cap was removed and CA1 was imaged using a miniscope mounted with an aluminum base plate while mice were under light anesthesia (*<*0.5% isoflurane) to allow the visualization of cell activity. When a satisfying field of view was found (large neuronal assembly, visible landmarks), the base plate was cemented above the GRIN lens, the miniscope was removed, and a plastic cap was placed on the base plate to protect the GRIN lens. For 0.5 mm GRIN lens LS and CA3 implants, the lens was glued to an aluminum baseplate prior to surgery. A 0.6 mm hole was drilled in the skull at the level of the injection site; an anchor screw was placed above the contralateral cerebellum to stabilize the implant. Baseplate-GRIN assemblies were attached to the miniscope and secured to the stereotaxic frame. While lowering the GRIN lens into the tissue, an increase in fluorescent signal indicates that the injection site has been reached. The GRIN lens and baseplate assembly was permanently secured using C&B-Metabond. Implants were targeted at the following coordinates: Anterior LS, AP: 0.86; ML: 0.35; DV: -2.9; Intermediate LS, AP: 0.38; ML: 0.35; DV: -2.65; Posterior LS, AP: 0.10; ML: 0.35; DV: -2.45; with DV *±* 0.2 to adjust for dorsal, ventral, or intermediate targets; CA3 implants were targeted at AP: -2.1; ML: 2.2; DV: -2.2. Three CA3 animals were only implanted with a 0.5 GRIN lens, and were baseplated 4 weeks later while under light anesthesia (*<*0.5% isofluorane). The baseplate was cemented after a satisfying field of view was found.

### In vivo behavioral procedures

#### Habituation

Mice were gently handled for ∼ 5 min over the course of one week, with progressive habituation to the miniscope attachment procedure. Animals were then water-scheduled or chow-scheduled (2 h access per day).

#### Miniscope recordings

Miniscopes (V3 and V4) were assembled using open-source instructions (miniscope.org). Imaging data were acquired using a CMOS imaging sensor (Aptina, MT9V032) and multiplexed through a lightweight coaxial cable. Data was acquired using a data acquisition (DAQ) box connected via a USB host controller (Cypress, CYUSB3013). Animal behavior was recorded using a consumer-grade webcam (Logitech) mounted above the recording setup. Calcium and behavioral data were recorded using miniscope.org open, source DAQ custom acquisition software. The DAQ simultaneously acquired behavioral and cellular imaging streams at 30 Hz as uncompressed .avi files of 1000 frames for 15 min recording sessions, along with corresponding timestamps to enable precise temporal alignment of behavioral and calcium imaging data. For open field recordings, the dimensions of the open fields were: 45 x 45 cm, 49 x 49, and 50 x 50, and were made of dark gray plexiglass. For linear track recordings, rewards (30 µl of 10% sucrose water) were placed at each end of either 100 or 134 cm linear tracks and the mouse had to consume one reward before getting the next one delivered. To assess the relative contribution of retrospective and prospective representations, one reward was replaced by a zone exposed to a pyroelectric sensor. The sensor instantly triggered a tone when triggered, and in turn triggered reward delivery on the opposite side. Finally, the sequential tone linear track task involved triggering pyroelectric sensors on either end of the track, each time triggering a new tone context (no tone, two rhythmic tones at distinct frequencies, and a continuous high-pitch tone which indicated that the reward had been delivered).

#### Post-mortem histological analyses

After completion of behavioral testing, mice were deeply anesthetized with a mixture of ketamine, xylazine, and acepromazide (100, 16, and 3 mg/kg, respectively, intraperitoneal injection) and perfused transcardially with 4% paraformaldehyde (PFA) in PBS. Brains were extracted and postfixed overnight in PFA at 4 °C and subsequently washed in PBS for an additional 24 h at 4°C. Brains and sections were cryoprotected in a solution of 30% ethylene glycol, 30% glycerol, and 40% PBS until used. Each brain was then sectioned at 50 *µ*m using a vibratome: every section was sequentially collected in 4 different 1.5 mL tubes to establish the GRIN lens location.

## Data analysis

### Automatized tracking of behavior

To extract information about the position, velocity, and head direction of mice, we used DeepLabCut [82, 83]. Briefly, we trained a model to detect mice’s ears, nose, body, and tail base. Head direction was estimated using the angle between either each ear or the nose and body, depending on measurement availability. Location data was interpolated to calcium imaging sampling frequency using linear interpolation. Velocity was extracted using equation (1).

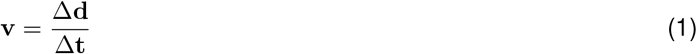

where **v** is the velocity, **d** is the distance and **t** time and subsequently smoothed the result by applying a Gaussian filter with *σ* = 33 ms to remove detection artifacts. Velocity signals were used to identify periods of locomotor activity (above 2 cm.s^−1^) and compute place, time, and distance modulation of activity specifically for those periods.

### Pre-processing and signal extraction

Calcium imaging data analysis was performed using Python 3.10.9. Video recordings were analyzed using the Minis-cope Analysis pipeline: https://github.com/etterguillaume/MiniscopeAnalysis Briefly, rigid (rotation and translation) motion correction was applied using NoRMCorre [84], and videos were spatially down-sampled 3*×* before concatenation. Calcium traces were extracted using CNMFe [43] using the following parameters:

- gSig = 3 pixels (width of Gaussian kernel)
- gSiz = 20 pixels (size of Gaussian kernel)
- background_model = ‘ring’
- spatial_algorithm = ‘hals’
- min_corr = 0.8 (minimum pixel correlation threshold)
- min_PNR = 8 (minimum peak-to-noise ratio threshold)

### Estimating neural activity

Raw calcium traces were filtered to remove high-frequency fluctuations and binarized: briefly, neurons were considered active when normalized calcium signal amplitude exceeded two standard deviations, and the first-order derivative was above 0 (see Etter *et al*. [47] for additional details on the methodology). To extract neurons tuning to specific variables, location, time, and distance were binned (location: 4 cm bins; time: 0.25 s bins; distance: 4 cm bins). From binarized signals, we computed the marginal likelihood of cells being active in a given behavioral state. Additionally, we computed Markovian transition probabilities to assess the tonic versus phasic nature of activity patterns.

### Information estimates

We estimated the information content of individual neurons using neural information (NI) between binarized activity *U* and discretized behavioral signal *V*, as defined by equation (2):

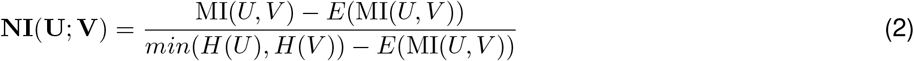

with *E* representing the expected mutual information (MI) between two random variables using a hypergeometric model of randomness, and MI defined by equation (3):

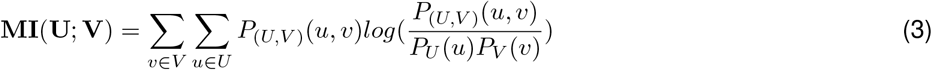

NI can be interpreted as the amount of uncertainty from a given behavioral variable that can be explained by neuronal activity, and is bound between 0 (no information) and 1 (one variable perfectly predicts the other). We show that this measure is corrected for low sampling, and insensitive to the number of bins used to discretize behavioral variables. The significance of independence between two variables is assessed using *χ*^2^ test and a p-value threshold of 0.05.

### Neural embeddings

Low-dimensional embeddings of raw calcium transients were learned using parametric UMAP with an autoencoder loss (pUMAP-AE; [46]), based on the original UMAP implementation [85]. Briefly, an autoencoder was trained with gradient descent and a dual objective function: (1) the encoder *f* was trained to minimize UMAP loss, and (2) the decoder *g* was trained to minimize reconstruction error such that *f* (*x*) = *z* and *g*(*z*) ≃ *x*, where x is the input neural activity and z the low-dimensional embedding with desirable properties such as smooth representation and preserved global structure. The the final loss function combines objectives (1) and (2). To evaluate our implementation of pUMAP-AE on neural data, we generate ground truth latent representations from which simulated neural data is sampled using a Poisson distribution [86].

#### Decoders

A k-Nearest Neighbors (kNN) regressor was trained to decode continuous behavioral variables (elapsed time, distance traveled, position, heading, velocity) from low-dimensional embedding. For categorical variables (e.g. tone, reward zone), a kNN classifier was used instead. To assess the decoding quality, we used the goodness of fit (*R*^2^) values. Additionally, we computed the median absolute error (MAE) between decoded values and ground truth for all variables but heading direction, for which we computed the absolute angle difference. Finally, a linear regression was used to learn a mapping between manifolds extracted from the train and test sets and the resulting goodness of fit (*R*^2^) was used as a proxy for representational stability.

#### Computational model

Place and grid cell rate maps were generated from a real exploration trajectory using the open source Python software RatInABox [87]. The activity rate of place cells were generated using equation (4):

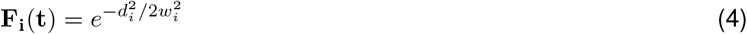

where *w* corresponds to the place cell width parameter which we set to 0.1 m, *i* is the neuron index, and *t* the timestamps.

Grid cell activity rates were generated as described previously [87]. Briefly, each grid cell was assigned preferred direction *θ*_*i*_, a random grid scale *λ*_*i*_ drawn from a Rayleigh distribution between 0.1 and 0.5 m), and a uniform random phase offset *ϕ*_*i*_ ∈ [0, 2*π*]. The firing rate of each grid cell is given by the thresholded sum of three cosines using equation (5):

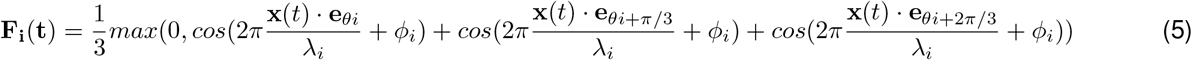

where *e*_*θ*_ is the unit vector pointing in the direction *θ*.

The respective activity rate from both cell types are then used to train a logistic regressor to predict the real activity of each individual neurons. To evaluate each model performance, we computed a F_1_ score for each predicted neuron using either a place or grid cell input model, which penalizes both incorrect classifications of active and inactive periods. We also computed a grid-over-place model fit ratio using equation 6:

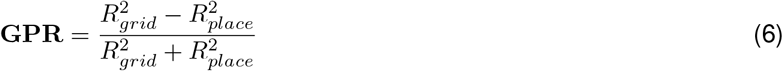

#### Train/test split

Low-dimensional embeddings were learned from 80% of the data, and tested on the held-out 20%. This latter test portion was further divided in 80% to train decoders and test accuracy on the remaining 20%.

#### Statistical analyses

Statistical analysis involved parametric tests when data distribution was normal and variance equal between groups, otherwise non-parametric tests were used. Where applicable, we assessed effect-size using Hedges **g** defined as:

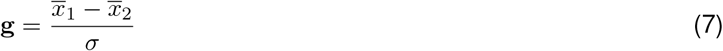

where 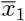 and 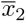 are the means of each group, and *σ* is the pooled standard deviation for both groups. Statistical tests and results are abbreviated as follows: 1-ANOVA, one-way ANOVA; 2-ANOVA, two-way ANOVA. ****, p < 0.0001; ***, p < 0.001; **, p < 0.01; *, p < 0.5; n.s., not significant.

## Data availability

The processed dataset generated in this study will be made publicly available upon publication on an Open Science Foundation repository or via a request to the corresponding authors.

## Code availability

All source codes used in the current study are available along with instructions here: https://github.com/etterguillaume/PyCaAn Extraction of calcium imaging data was done using: https://github.com/etterguillaume/MiniscopeAnalysis(v1.0) This extraction pipeline leverages motion correction from NoRMCorre: https://github.com/flatironinstitute/NoRMCorre (v0.1.1) as well as CNMFe video processing: https://github.com/zhoupc/CNMF_E (v1.1.2) specifically for V3 miniscopes. Computational models leveraged RatInABox [87]: https://github.com/RatInABox-Lab/RatInABox (v1.10.0)

## Acknowledgments

This work was supported by funding from the Canadian Institutes for Health Research (CIHR) Foundation Program FDN-148478, the Natural Sciences and Engineering Research Council of Canada (NSERC) Discovery Grant RGPIN-2020-06717, and a Tier 1 Canada Research Chair to S.W. S.V.D.V. was supported by a Vanier Canada Graduate Scholarship and the Richard H. Tomlinson Doctoral Fellowship. C-AM was supported by a FRQS postdoctoral fellowship. The funders had no role in study design, data collection and analysis, decision to publish, or preparation of the paper.

## Author contributions

Conceptualization: GE

Methodology: GE

Investigation: GE, SVDV, CAM

Visualization: GE

Funding acquisition: SW

Project administration: SW

Supervision: SW, GE

Writing – original draft: GE

Writing – review and editing: GE, SVDV, CAM, MEH, SW

## Supplementary information

**Supplementary figure 1:**
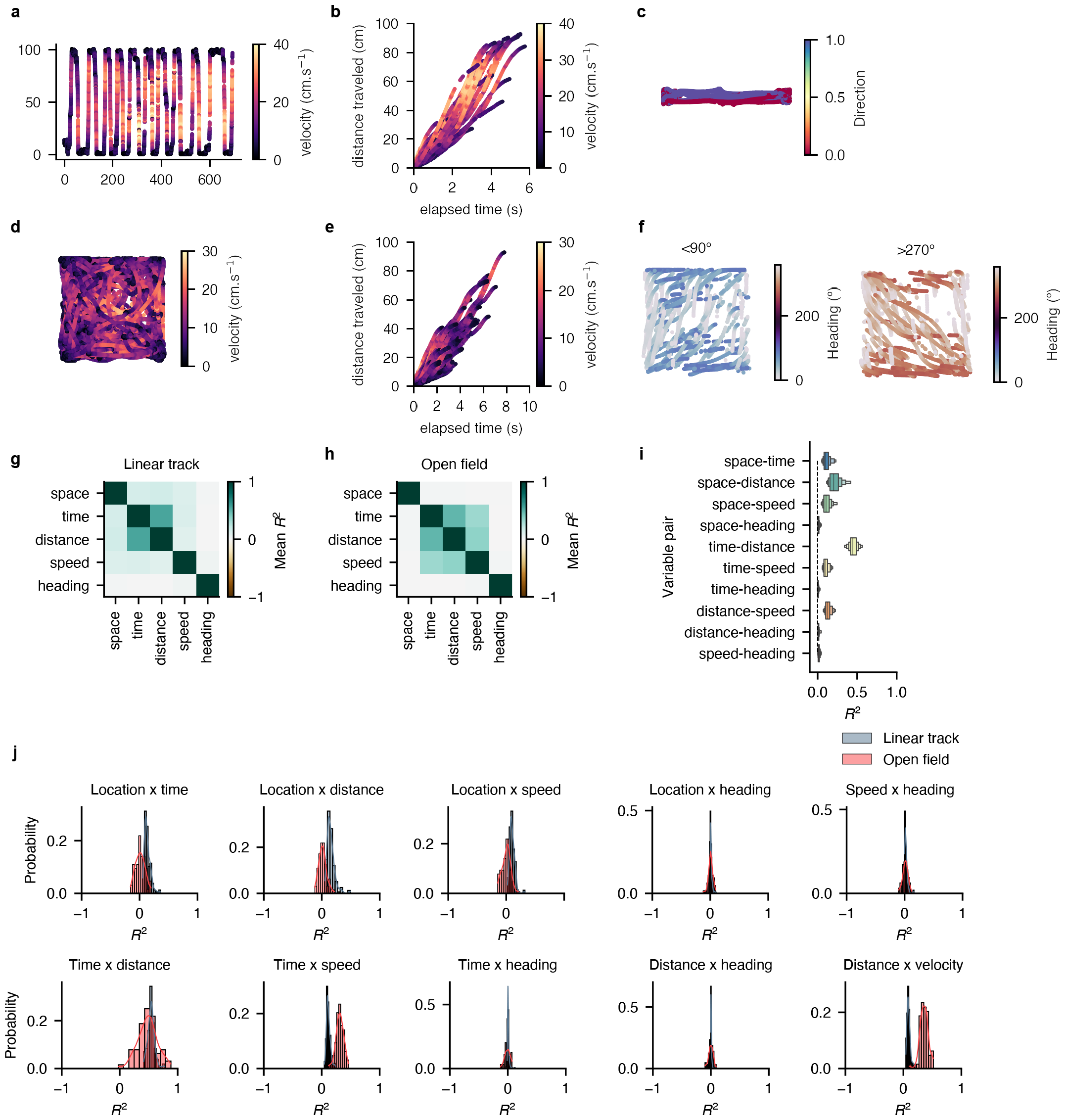
Spatiotemporal covariates are separable. **a**, example trajectory on the linear track. **b**, example comparison between elapsed time, distance traveled, and speed on the linear track. **c**, example direction of travel (left vs right) on the linear track. **d**, example trajectory in the open field. Color indicates speed. **e**, example trajectories for a given direction (<90º). **f**, sample but for directions above 270º. **g**, mean neural information values between behavioral covariates on the linear track. **h** same for open field recordings. **i**, summary of neural information values between behavioral variables (linear and open mazes are pooled here). **j**, histogram distribution for neural information values between pairs of behavioral variables. Blue, linear track; red, open field.

**Supplementary figure 2:**
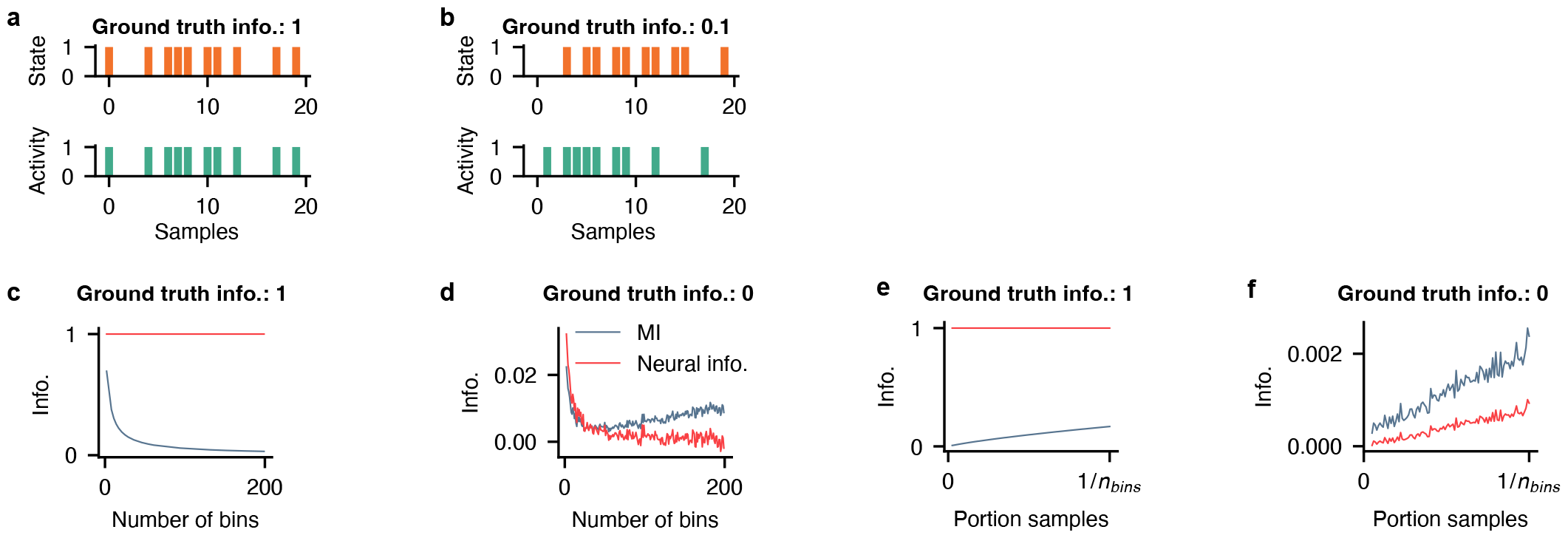
Unbiased extraction of neural information. **a**, simulated neural activity (green) and binary state (orange) where the ground truth information between the two variables is set to 1. **b**, same but for a scenario where ground truth information is set to 0.1. **c**, neural information (red) and mutual information (MI, blue) as a function of the number of bins used to discretize the state variable, in conditions where the ground truth information between simulated neural activity and states is set to 1. **d**, same for scenarios where the ground truth information between the two variables is set to 0. **e**, neural information (red) and mutual information (MI, blue) as a function of the portion of data sampled to compute both metrics, in conditions where the ground truth information between simulated neural activity and states is set to 1. **f**, same for scenarios where the ground truth information between the two variables is set to 0.

**Supplementary figure 3:**
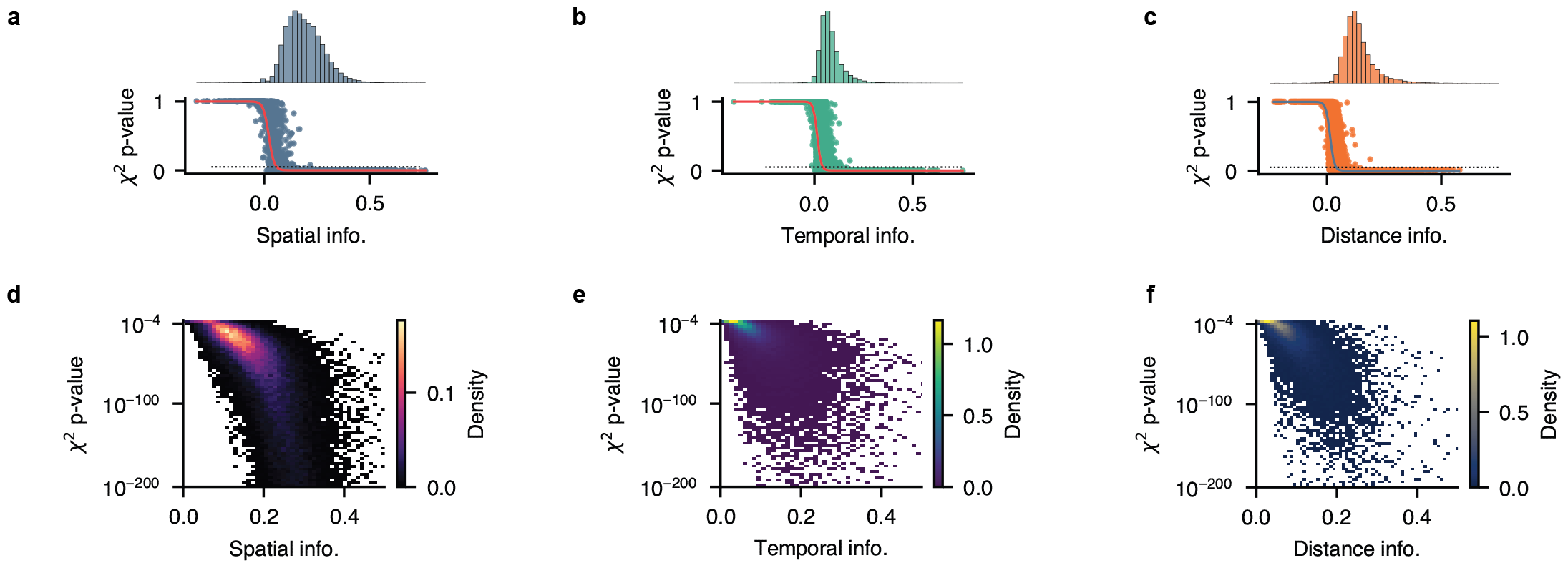
Distribution and significance of spatial and temporal information. **a-c**, distribution of neural information against *χ*^2^ p-value for space (**a**), elapsed time (**b**), and distance traveled (**c**). **d-f**, same but where the density of data points is displayed instead of individual points, and the vertical axis is shown on a log scale.

**Supplementary figure 4:**
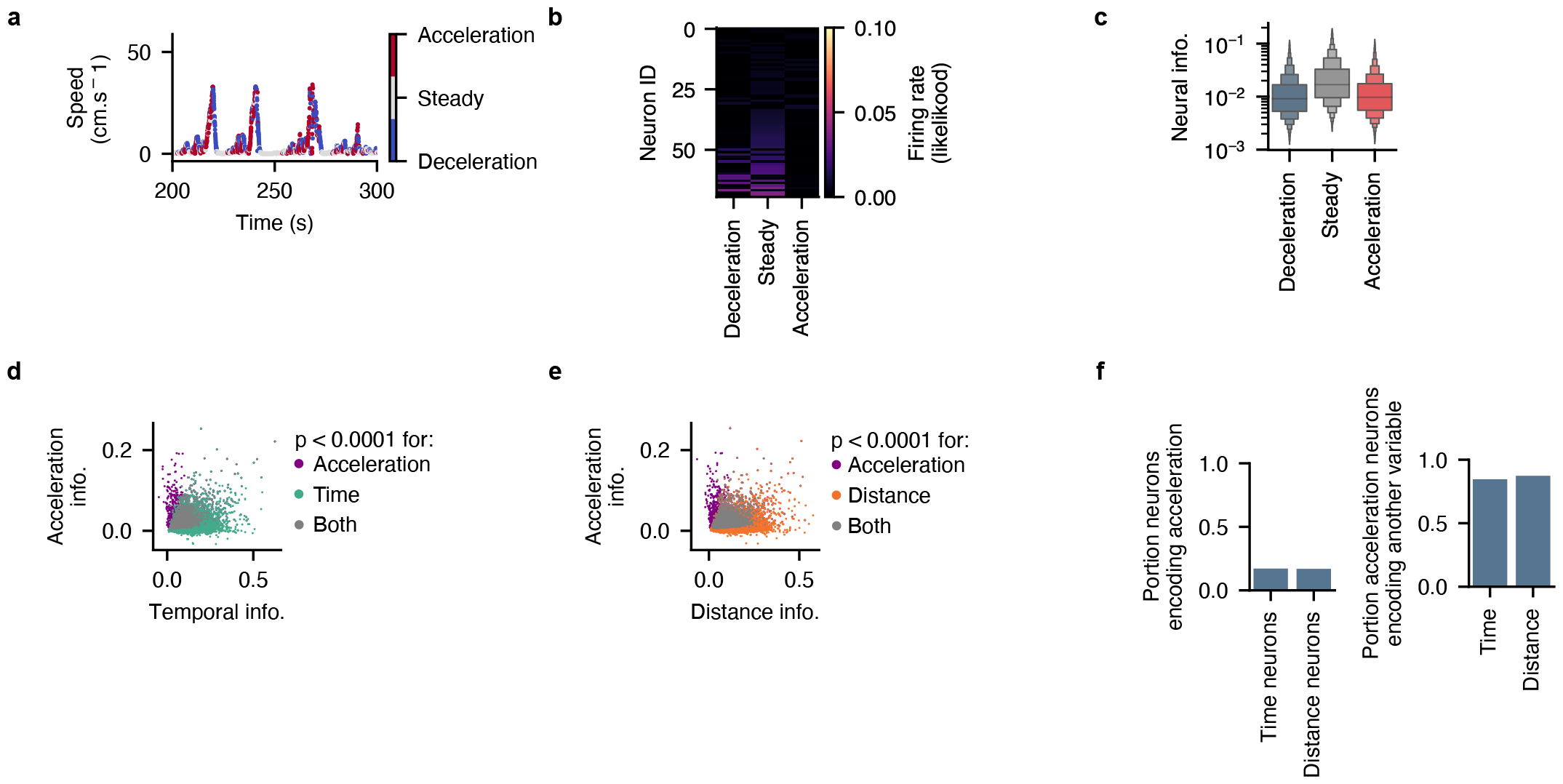
Most time and distance cells do not encode acceleration. **a**, periods of acceleration, steady speed, and deceleration from an example mouse. **b**, neural information for each state for a subset of example neurons. **c**, average information for each acceleration state. **d**, temporal and acceleration information for neurons pooled from n = 19 independent mice. Each point represents a neuron, color coded for significance for acceleration, time, or both. **e**, same but for distance and acceleration. **f**, portion of time and distance neurons also encoding acceleration (left). Right, portion of acceleration neurons also encoding time or distance.

**Supplementary figure 5:**
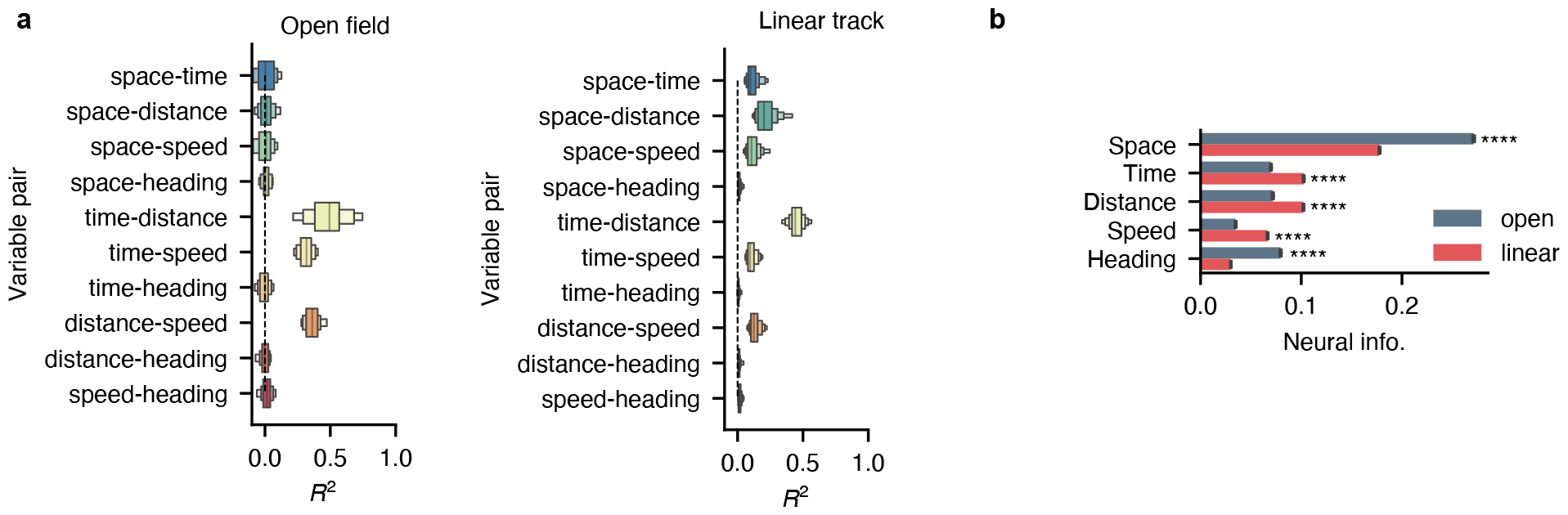
Extracting neural representations of time and distance in open environments. **a**, example locomotor events in an open field (top) and the relationship between elapsed time and distance traveled for all trajectories in a representative session. **b**, correlation coefficient for each pair of behavioral variable in the open field. **c**, example rate map in the open field for an example CA1 neuron that significantly encodes space. Yellow, maximum firing rate that corresponds to the number in the bottom left corner; dark blue, no activity. **d**, neural information for allocentric and idiothetic variables in open (blue) versus linear environments (red).

**Supplementary figure 6:**
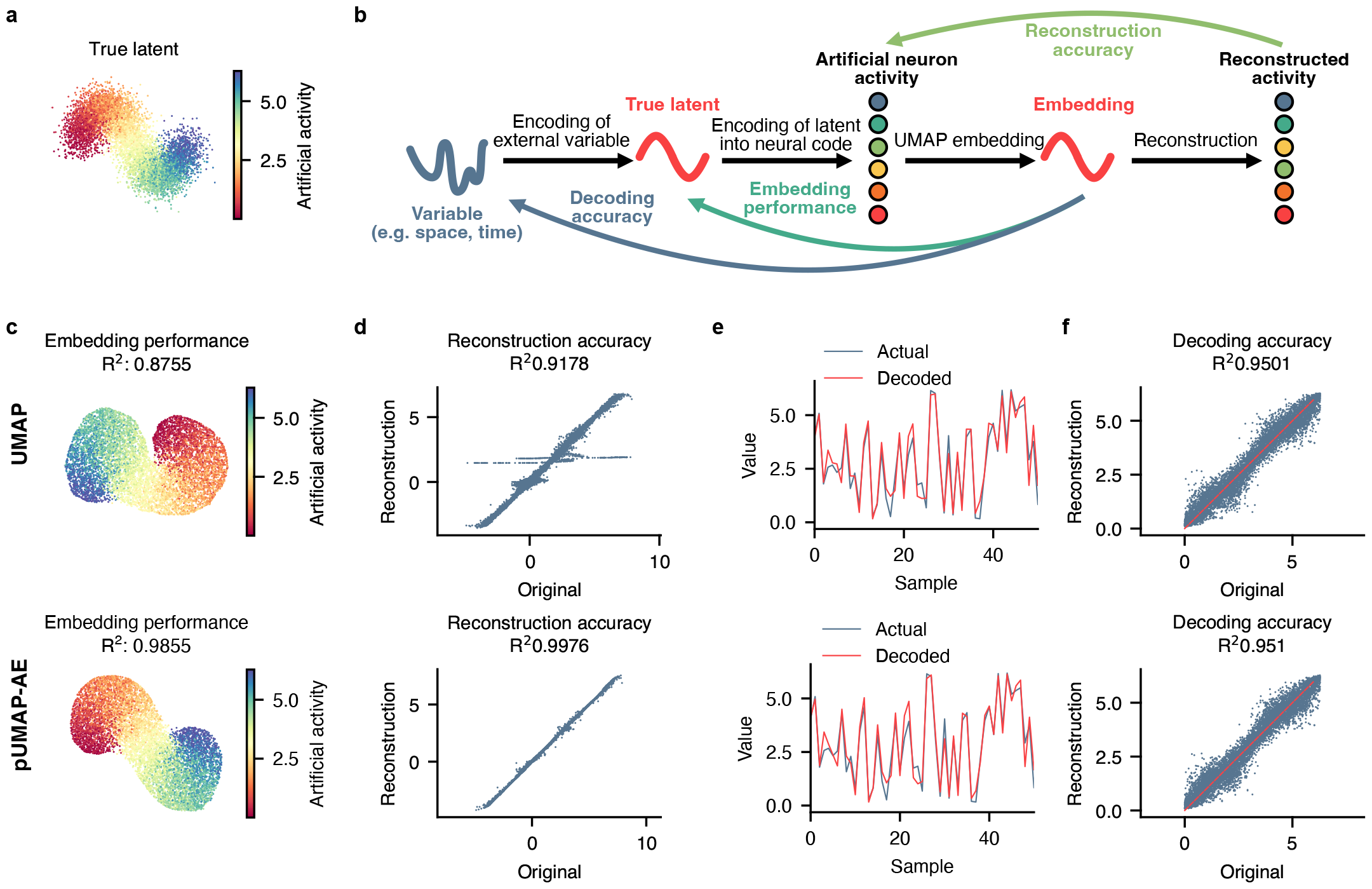
Embedding performance on simulated neural activity. **a**, simulated latent used to generate artificial activity. **b**, given a known label and latent, artificial neural activities are generated using Poisson sampling. **c-f**, performance comparison between vanilla UMAP (top) and parametric UMAP with autoencoder loss (bottom). **c**, learned latent and associated embedding performance, as assessed using a linear regression between the true and estimated latent representation. **d**, original versus reconstructed activity and associated reconstruction accuracy. **e**, actual variable value at the origin of the true latent versus variable value decoded from the embedding. **f**, decoding accuracy.

**Supplementary figure 7:**
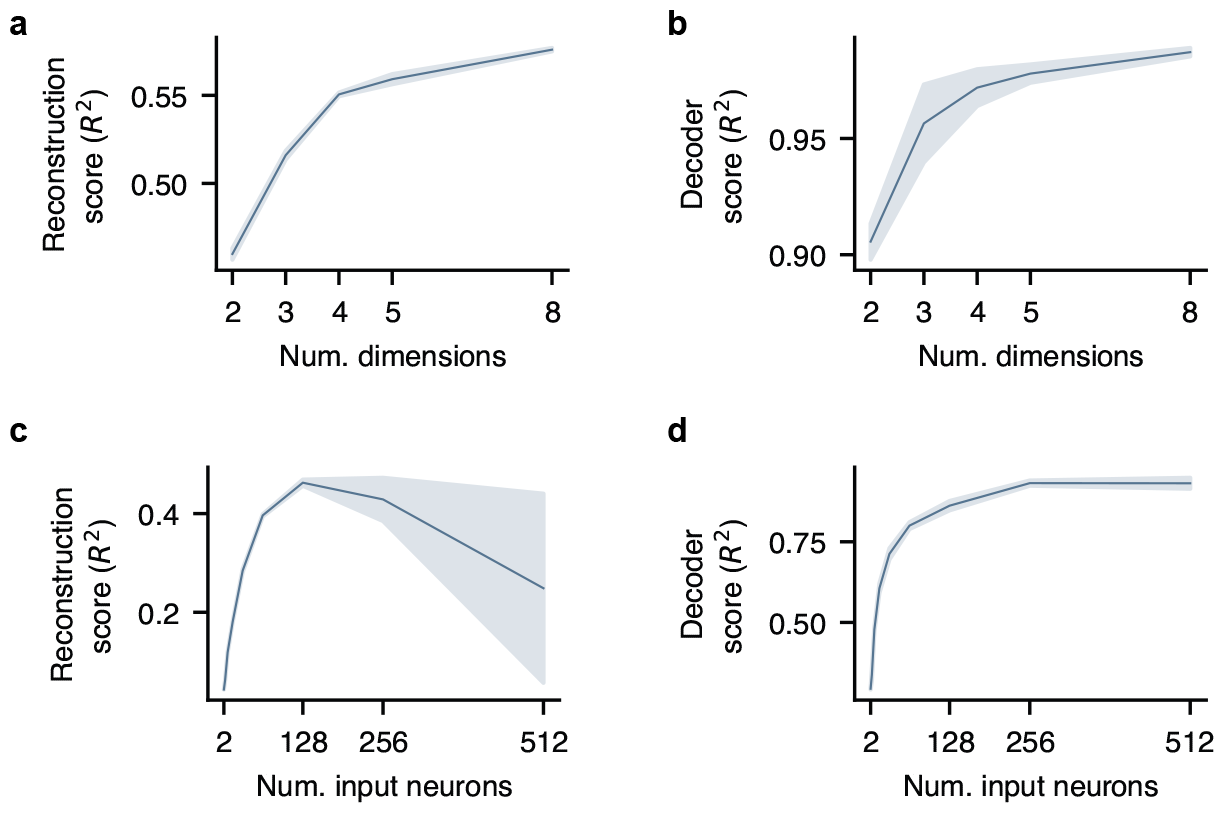
Parameter optimization for low-dimensional embedding. **a**, reconstruction score (*R*^2^) using 2, 3, 4, 5, or 8 embedding dimensions. **b**, decoding score (*R*^2^) using 2, 3, 4, 5, or 8 embedding dimensions. **c**, reconstruction score (*R*^2^) using 2, 4, 8, 16, 32, 64, 128, 256, or 512 input neurons. **d**, decoding score (*R*^2^) using 2, 4, 8, 16, 32, 64, 128, 256, or 512 input neurons.

**Supplementary figure 8:**
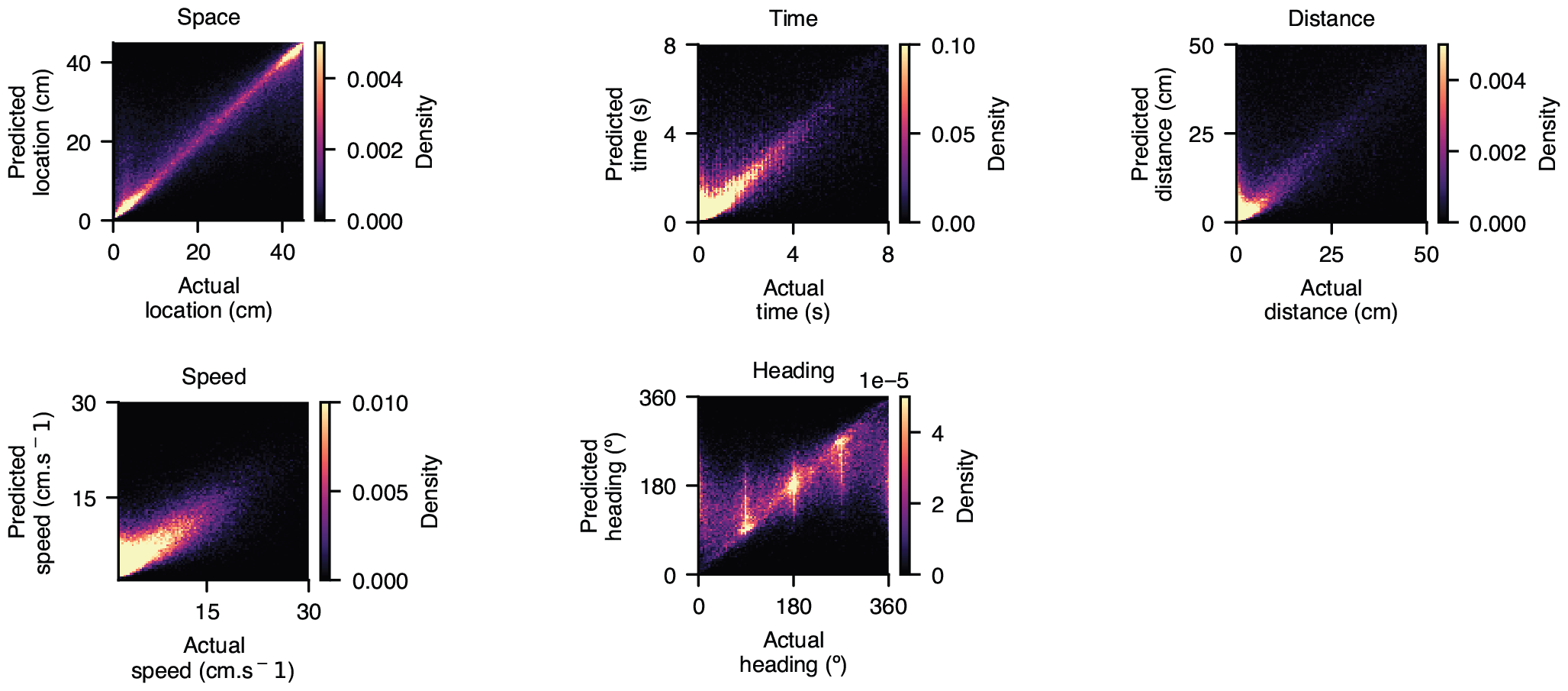
Decoding confusion matrices. Decoding confusion matrices for each modality, using n = 100 quantization bins.

